# Fur4 mediated uracil-scavenging to screen for surface protein regulators

**DOI:** 10.1101/2021.05.27.445995

**Authors:** Katherine M. Paine, Gabrielle B. Ecclestone, Chris MacDonald

## Abstract

Cell surface membrane proteins perform diverse and critical functions and are spatially and temporally regulated by membrane trafficking pathways. Although perturbations in these pathways underlie many pathologies, our understanding of these pathways at a mechanistic level remains incomplete. Using yeast as a model, we have developed an assay that reports on the surface activity of the Fur4 uracil permease in uracil auxotroph strains grown in the presence of limited uracil. This assay was used to screen a haploid deletion library that identified mutants with both diminished and enhanced comparative growth in restricted uracil media. Factors identified, including various multi-subunit complexes, were enriched for membrane trafficking and transcriptional functions, in addition to various uncharacterised genes. Bioinformatic analysis of expression profiles from many strains lacking identified transcription factors required for efficient uracil-scavenging revealed they control expression of other uracil-scavenging factors, in addition to membrane trafficking genes essential for viability, and therefore not represented in the screen. Finally, we performed a secondary mating factor secretion screen to functionally categorise factors implicated in uracil-scavenging, most of which are conserved throughout evolution.

## INTRODUCTION

Cell surface membrane proteins are regulated by a variety of overlapping and often co-regulated membrane trafficking pathways. Surface cargoes are co-translationally imported into the endoplasmic reticulum (ER)^1^ before transiting the secretory pathway to the plasma membrane (PM) for function^2^. Surface proteins are internalised via clathrin mediated endocytosis, followed by recycling back to the PM or entering the lysosomal degradation pathway^3^. These pathways allow surface localisation and activity of myriad proteins to be precisely controlled to meet cellular demands, for example during the cell cycle or in response to reduced nutrient availability; however, these pathways remain incompletely characterised. The budding yeast system has been useful to discover and define membrane trafficking mechanisms. Yeast cells uptake nutrients from their external environment by a variety of transporters that localise to the PM, such as transporters for sugars, metal ions and vitamins^4–6^. Amino acids and nucleosides are also actively imported via nutrient transporters, for example Gap1 broadly uptakes amino acids^7^, Mup1 uptakes methionine^8^ and Fur4 uptakes uracil^9^. Permease activity can be controlled by changes in transporter expression and the rate of turnover by ubiquitin mediated vacuolar degradation, in addition to spatiotemporal control between eisosomes and other regions of the PM^10–13^. To develop an assay that reports on nutrient uptake via surface transporters, we focussed on the uracil permease Fur4^9^, which is controlled by the above-described trafficking pathways and regulatory mechanisms. For example, the presence of uracil downregulates expression of *FUR4*^14,15^ whilst also triggering endocytosis and Rsp5-mediated ubiquitination and degradation of Fur4^16,17^. Furthermore, Fur4 activity is regulated in response to metabolic stress via storage in eisosomes^18–20^. Fur4-mediated uptake of uracil might be considered particularly important for many lab strains that cannot synthesise uracil biosynthetically, due to disruption of the orotidine-5’-phosphate decarboxylase *URA3* gene (e.g. *ura3-52* or *ura3*Δ), which impose the useful selection characteristics of auxotrophy and resistance to 5-Fluoroorotic Acid^21^. Indeed, many genome-wide libraries^22–28^ have been created from designer parental *ura3*Δ strains^29^. We therefore chose Fur4-mediated uptake to develop a simple and cost-effective growth assay that indirectly reports on the surface-mediated uptake of uracil, which we have used to screen a haploid library of deletion mutants for factors that regulate Fur4 trafficking, many of which are conserved across evolution.

## RESULTS & DISCUSSION

### Uracil-scavenging yeast growth assay

The uracil permease Fur4 is dispensable for growth in rich media but is critically required when uracil-auxotroph cells are grown in synthetic defined media containing replete (4 mg/L) uracil (**Figure 1A - 1C**). Importantly, the robust Fur4-dependent growth in BY4742 cells, which harbour a *ura3*Δ mutation^29^ herein referred to as wild-type, corresponds to the concentration of available uracil. There is a significantly reduced rate of growth when wild-type cells are grown in media containing only 0.1 mg/L uracil, but Fur4 dependent uracil-scavenging supports growth (**Figure 1B, 1C**). Fur4 has been previously shown to respond to extracellular uracil^14^, and we confirm steady state surface localisation of Fur4 tagged with mNeonGreen (mNG) is redistributed to FM4-64 stained vacuoles following 1-hour of uracil addition to the media (**Figure 1D**). Collectively these results show that the activity of the uracil sensitive permease Fur4 correlates with cellular growth in limited uracil conditions. To test if low uracil-specific growth could be used to screen for membrane trafficking factors that influence Fur4 surface levels, we next compared Fur4-mNG localisation in wild-type cells and mutants that mis-localise Fur4 (**Figure 2A**). As expected, a temperature sensitive *sec7-1* allele, which disrupts activity of the Sec7 Arf-exchange factor required for transit through the Golgi^30,31^, inhibits trafficking of Fur4 through the secretory pathway with reduced levels at the PM. Fur4-mNG localisation is also affected in *vps2*Δ ESCRT mutants that do not permit vacuolar sorting^32^, where Fur4-mNG instead accumulates in endosomes. However, uracil-scavenging is likely efficient in *vps2*Δ cells as significant Fur4-mNG recycling back to the PM is observed, unlike Mup1-GFP that is trapped by Snf7-oligomers^33^. In contrast, surface levels of Fur4-mNG was greatly reduced in both *rcy1*Δ or *nhx1*Δ mutants, (**Figure 2A**), which lack factors required for efficient endosomal recycling^34–36^. Therefore, we compared growth of *rcy1*Δ and *nhx1*Δ mutants with wild-type cells on plates of varying uracil concentrations. There was no statistically significant difference in growth between wild-type and trafficking mutant cells in the range of 1 mg/L - 32 mg/L uracil, but at lower uracil concentrations the *rcy1*Δ and *nhx1*Δ mutants, which have reduced surface Fur4, exhibit a low-uracil specific growth defect (**Figure 2B, 2C**). Although significant defects were observed at 0.5 mg/L and 0.25 mg/L uracil, we selected 0.1 mg/L and 0.05 mg/L uracil for optimal resolution scavenging conditions, as wild-type cells grow efficiently but *rcy1*Δ or *nhx1*Δ both show dramatically reduced growth. We tested this concept with another surface-localised nutrient transporter, the methionine transporter Mup1 grown in methionine auxotroph (*met15*Δ) cells but no methionine concentration that supports growth could distinguish trafficking mutants (**Figure S1**). We assume differences in steady state surface levels, substrate affinity and uptake pathways^8^ account for this.

**FIGURE 1:**
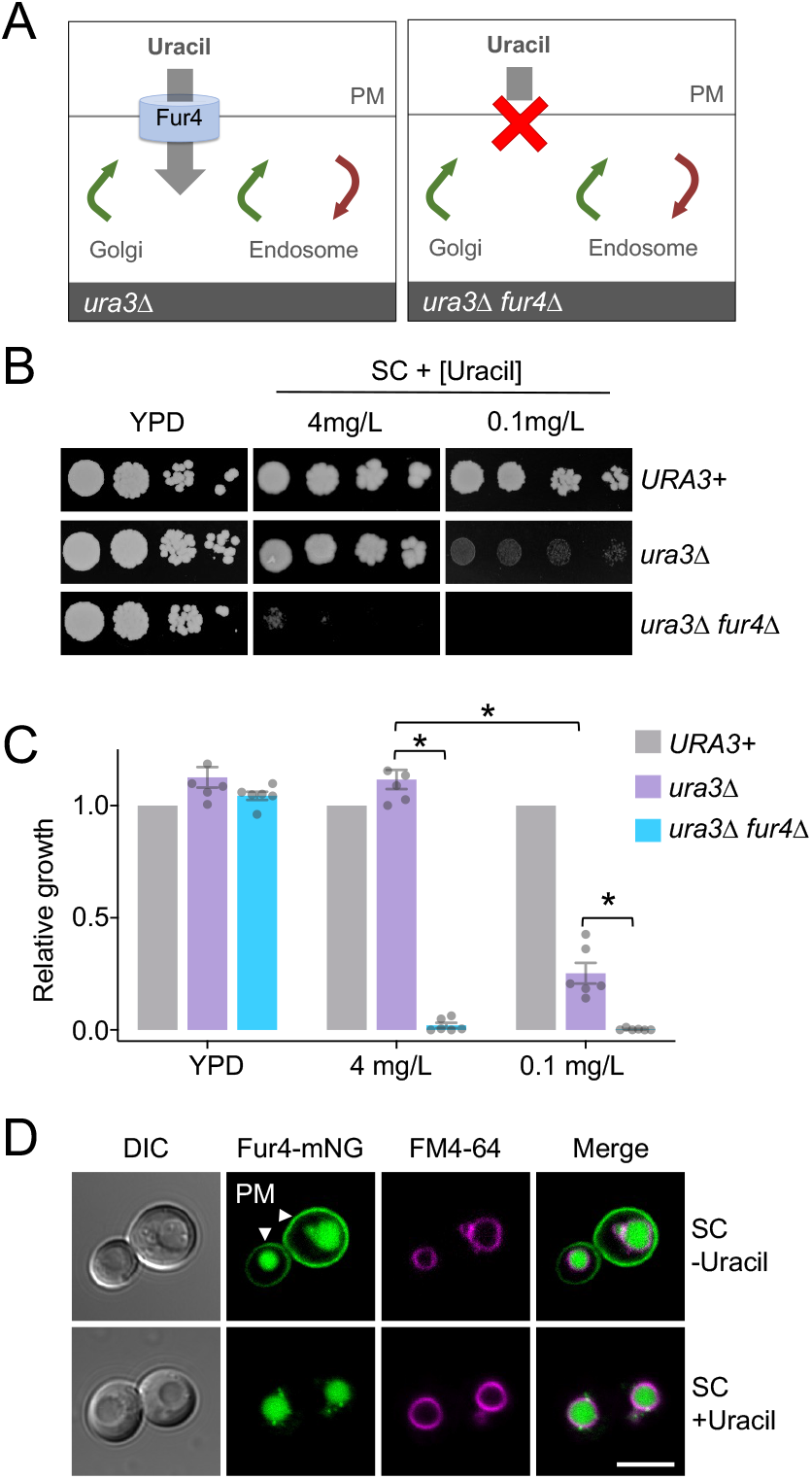
Low uracil growth relies on the Fur4 transporter. *A:* Schematic illustration showing uracil-auxotroph cells (*ura3*Δ) with (left) and without (right) the uracil permease Fur4. *B:* Indicated yeast strains were grown to mid-log phase before plating on rich (YPD) and synthetic complete (SC) media containing either 4 mg/L or 0.1 mg/L uracil. *C:* Quantification of yeast growth from *B*, asterisks (*) indicate Student’s *t*-test comparisons p < 0.001. *D:* Wild-type cells expressing Fur4-mNeonGreen (Fur4-mNG) from its endogenous promoter were labeled with FM4-64 for 1-hour, grown to mid-log phase in SC-Ura media (upper) or in the presence of 40 µg/ml uracil for 1 hour (lower) prior to confocal microscopy. White arrow heads indicate plasma membrane (PM) signal. Scale bar = 5µm.

**FIGURE 2:**
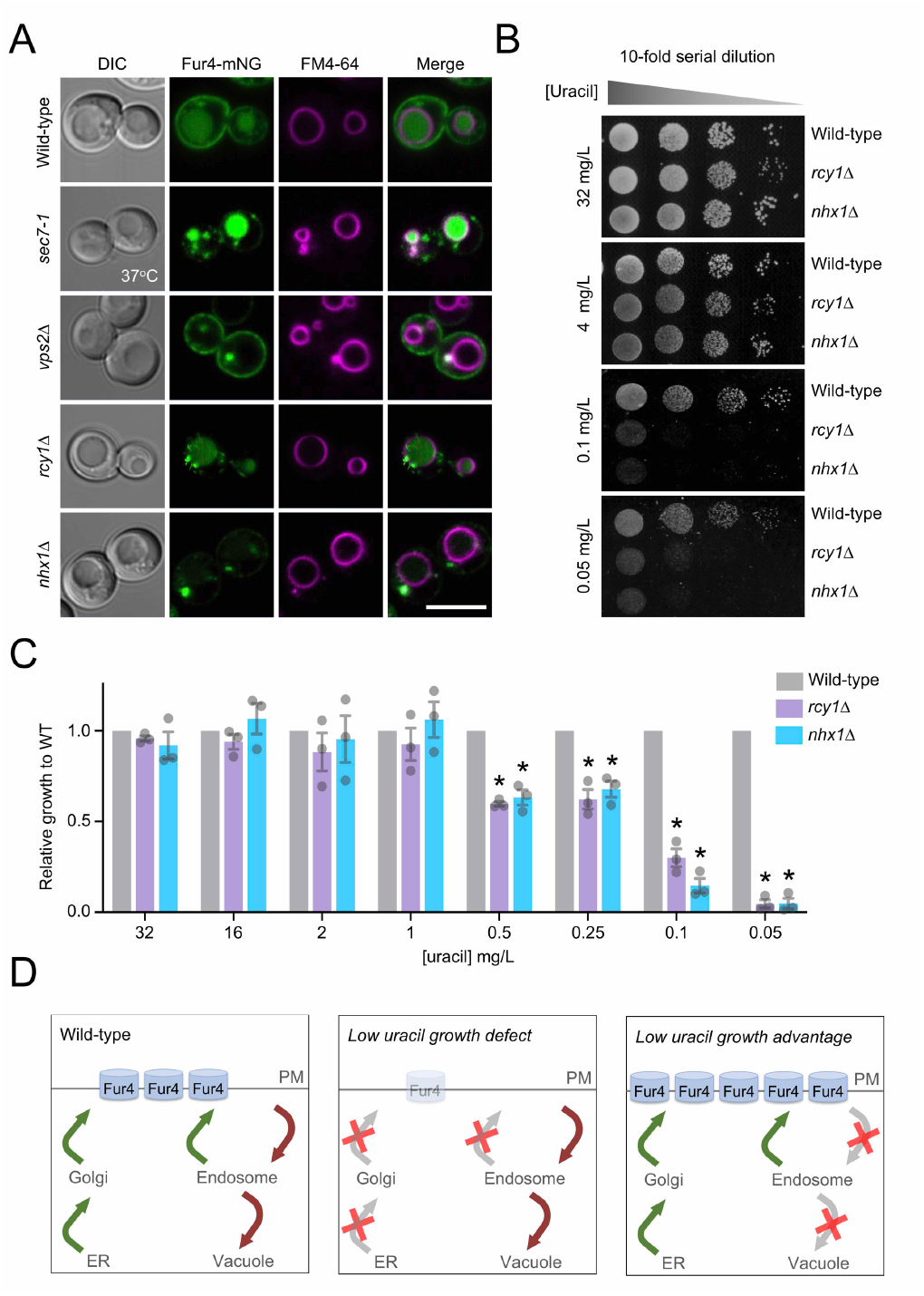
Surface localization of Fur4 is required for growth in low-uracil. *A:* Indicated strains expressing Fur4-mNeonGreen (mNG) were labelled with a 1-hour FM4-64 pulse prior to growth in label-free chase media for 2 hours and preparation for confocal imaging. Scale bar = 5 µm. *B:* Wild-type, *rcy1*Δ and *nhx1*Δ cells grown to mid-log phase were spotted in a 10-fold serial dilution onto plates titrated with indicated concentrations of uracil and grown at 30°C for 3 days. *C:* Growth of strains from *B* were quantified, asterisks (*) indicate Student’s *t*-test comparisons of mutants with wild-type cells, p < 0.005. *D:* Schematic diagrams showing the predicted effects on Fur4 following different trafficking pathway perturbations.

### A genetic screen for uracil-scavenging mutants

We hypothesised that mutants with growth similar to wild-type in replete uracil but differences specifically at low uracil could be used to identify mutants from a non-essential haploid deletion library^22^ with perturbed surface levels of Fur4 (**Figure 2D**). Cultured yeast strains representing 5132 different mutants were diluted and spotted out (16 replicates of each) on to solid agar media containing replete (4 mg/L) and limited (0.1 mg/L and 0.05 mg/L) uracil concentrations (**Figure 3A**). As expected, mutants with growth differences in uracil-replete media were observed, but the screen was specifically focused on differences in growth between high and low uracil concentrations. This is exemplified by the mutants used to calibrate the assay, *rcy1*Δ and *nhx1*Δ, which both exhibit growth defects specifically in low-uracil when compared with neighboring mutants (**Figure 3B**). A low stringency scoring system was used to identify 208 null mutants for follow up analysis. Candidates were grown to mid-log phase then serially diluted and spotted on 4 mg/L, 0.1 mg/L and 0.05 mg/L uracil media. Growth was quantified for the 208 mutant strains compared to a wild-type control from the same plate, and then values were used to compare growth across uracil concentrations (**Table S1**). Statistical comparisons of mutants from these optimized growth assays revealed 58 mutants, such as *cla4*Δ (**Figure 3C**), that did not show a significant difference in growth compared to wild-type. However, 126 mutants with growth defects specifically in uracil-scavenging conditions were identified (**Figure 3D**), ranging from relatively subtle defects (e.g. *vps74*Δ at 0.1 mg/L = 0.78 ± 0.04) to extreme (e.g. *vps3*Δ at 0.1 mg/L = 0.004 ± 0.003). Furthermore, although the assay was calibrated for mutants with defective growth in low-uracil, 24 mutants with enhanced growth compared to wild-type were validated. We note many of these strains exhibit growth defects in uracil replete media, such as *ipk1*Δ (**Figure 3C**), allowing for benefits to be observed at low uracil (**Table S1**).

**FIGURE 3:**
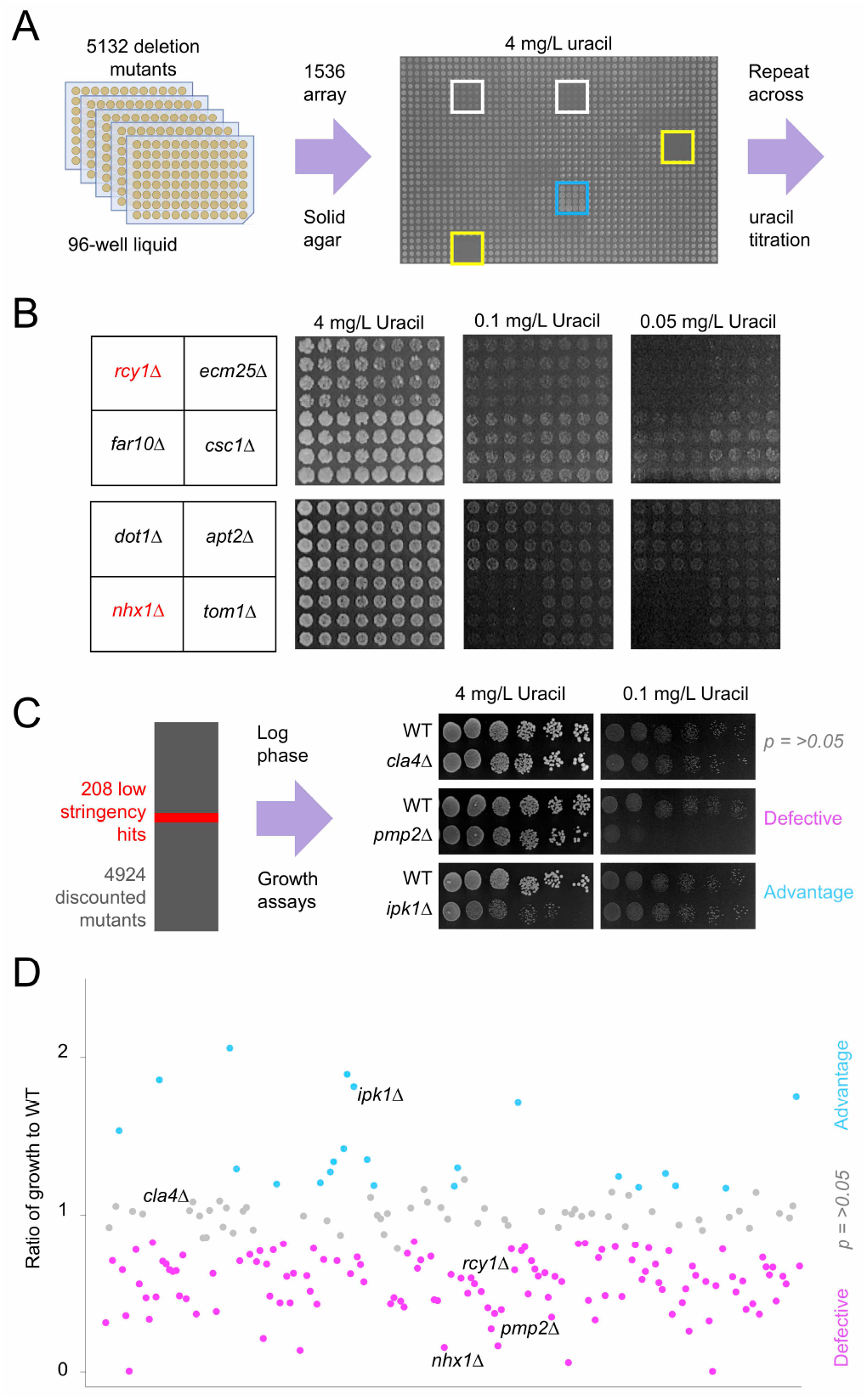
A genetic screen for mutants that affect uracil-scavenging. *A:* Yeast strains grown overnight in 96-well plates were diluted 20-fold in water then replicated 16 times onto solid agar plates containing varying uracil concentrations. An example 4 mg/L uracil plate shows identifier wells (yellow), alongside strains with defective (white) and accelerated (blue) growth. *B:* Example of screen data showing indicated mutants grown on media containing 4 mg/L, 0.1 mg/L and 0.05 mg/L uracil, including known Fur4 trafficking mutants (red). *C:* The screen identified 208 candidates that were subsequently grown to mid-log phase and spotted out over a 6-step, 10-fold serial dilution onto high (4 mg/L) and lower (0.1 mg/L and 0.05 mg/L) uracil containing media. Uracil-related cellular growth relative to wild-type was quantified and categorized as defective (e.g. *pmp2*Δ), advantageous (e.g. *ipk1*Δ) or not significantly altered (e.g. *cla4*Δ). *D:* Ratio of relative growth between 4 mg/L and 0.1 mg/L uracil from *C* was plotted for all candidates.

### Screen enriched for molecular complexes

A comparative orthologue search^37^ of uracil-scavenging mutants identified 94 highly conserved genes, corresponding to 267 human orthologues associated with 87 distinct diseases (**Table S2**). Gene Ontology (GO) enrichments were performed^38^ to revealed a number of cellular component enrichments of molecular complexes (**Figure S2**), including the GET complex and the prefoldin complex (**Figure 4A**). The identification of multiple complex members suggests the screen was stringent and robust. For example, deletion of *GET1, GET2*, or *GET3* results in low-uracil growth defects (**Figure 4B**). As the GET complex is required for sorting of tail-anchored single-pass membrane proteins^39^, many of which are important factors in secretory pathway trafficking^40^, we presume the role of the GET complex in efficient surface trafficking of Fur4 is via an indirect membrane trafficking mechanism. Various prefoldin complex members were also identified as having defective growth specifically on low uracil (**Figure 4C**), implicating it as a potential regulator of Fur4 trafficking. This could be explained by prefoldin-mediated assembly of cytoskeleton proteins^41,42^. Indeed, actin filament structures observed in wild-type cells are absent in prefoldin mutant cells *pfd1*Δ and *pac10*Δ (**Figure 4D**), indicating impaired microtubule function that could adversely affect correct trafficking of Fur4 to the surface. However, the prefoldin complex is also involved in transcriptional elongation^43^, so its role in uracil-scavenging could be indirect.

**Figure 4:**
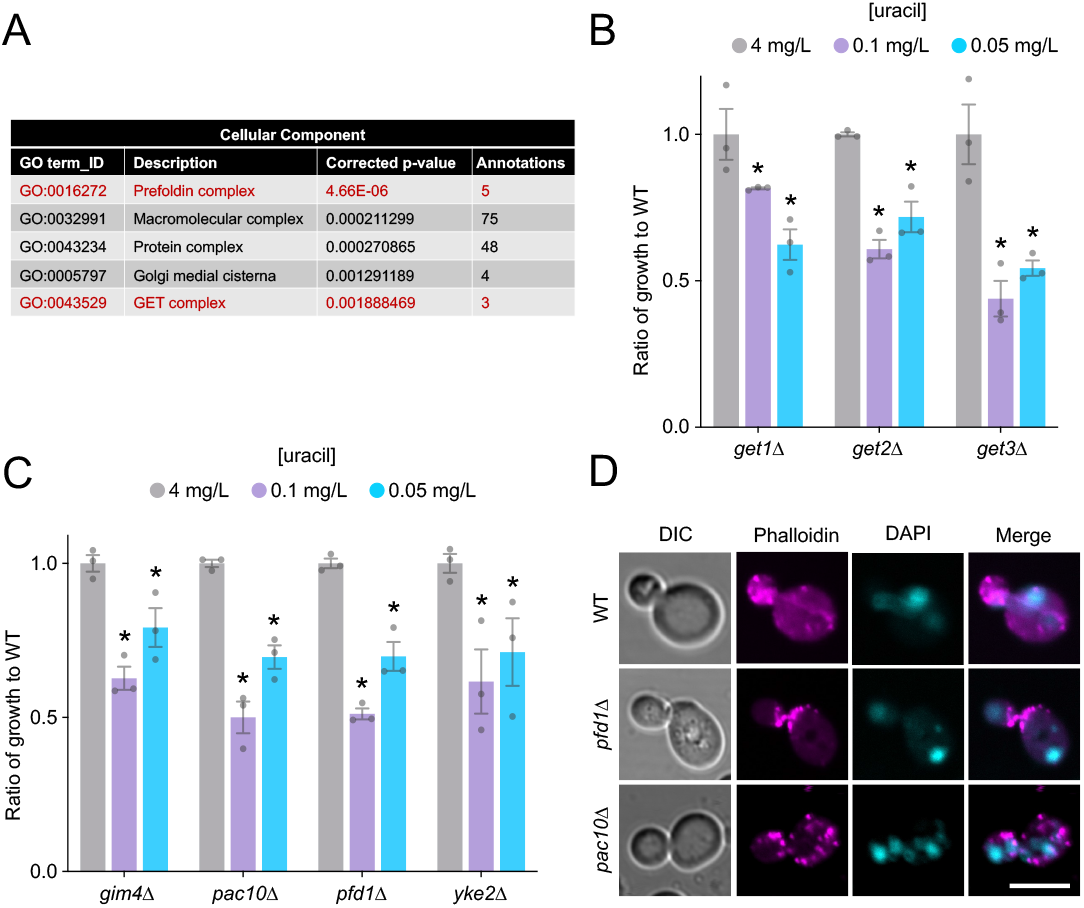
Uracil-scavenging screen identifies multi-subunit complexes. *A:* Gene Ontology enrichment analysis for cellular component of the 150 factors identified from the screen. *B:* Ratio of growth compared to wild-type (WT) cells at 4 mg/L, 0.1 mg/L and 0.05 mg/L uracil for GET complex mutants: *get1Δ, get2*Δ and *get3*Δ. Asterisks (*) indicate Student’s *t*-test comparisons p < 0.001. *C:* Ratio of growth compared to wild-type (WT) cells at 4 mg/L, 0.1 mg/L and 0.05 mg/L uracil for indicated Prefoldin complex mutants: *gim4Δ, pac10Δ, pfd1*Δ and *yke2*Δ. Asterisks (*) indicate Student’s *t*-test comparisons p < 0.01. *D:* Confocal microscopy of wild-type (WT) and prefoldin mutants fixed with 4% paraformaldehyde and stained with Phalloidin-594 and DAPI. Scale bar = 5µm.

### Screen enriched for trafficking and transcriptional machinery

GO enrichments for biological process revealed almost a third (44/150) of annotations were for machinery associated with membrane trafficking and signalling (**Figure 5A, Table S3**), including the blind identification of *rcy1*Δ and *nhx1*Δ mutants that were used to calibrate the assay (**Figure 2**). The other biological process significantly enriched was transcription, including 3 prefoldin subunits annotations (**Figure 5A, Table S3**). Physically interacting transcription factors were identified (**Figure S2**), such as members of the COMPASS complex, Swd1, Swd3, and Sdc1^44^ alongside the Swi3/Snf5 pair^45^ amongst others, all of which exhibit significant defects in low uracil (**Figure 5B**). We reasoned transcriptional regulators could be indirectly involved in Fur4 membrane trafficking, controlling gene expression of either essential genes not tested in the primary screen or mutants identified from the screen itself. To explore transcription factors (TFs) implicated in uracil-scavenging, we assembled genome-wide expression datasets for wild-type cells versus 28 TF-null mutants from a large-scale microarray analysis^46^. A matrix of all mutants showed high correlation of associated factors, such as known complex members (**Figure S3**). Cross-referencing expression data for 1183 genes that are essential for viability (**Table S4**), followed by hierarchical clustering was used to generate a heat map of related gene expression changes (**Figure 5C**). Strains such as *ies2Δ, bre1*Δ and *uba4*Δ created distinct expression signatures, but others were similar, such as each of the COMPASS complex mutants *swd1Δ, swd3*Δ, and *sdc1*Δ. Particularly modulated clusters of genes were identified from this analysis (**Table S5**), including genes associated with membrane trafficking that we chose for experimental testing (Pma1, Gpi8, Mrs6, Gpi12 and Sec62). To achieve this, we performed uracil-scavenging assays with strains containing Decreased Abundance by mRNA Perturbation (DAmP) cassettes at the 3’ UTR of each candidate^47^. This analysis revealed *sec62-DAmP* cells, which have very low protein levels of Sec62^48^, have specific growth defects in low uracil (**Figure 5D**). *SEC62* expression was greatly reduced upon deletion of several TFs from the screen, for example *uba4*Δ and *met18*Δ mutants (**Figure 5E**). The uracil-scavenging defects in *sec62-DAmP* cells can be explained in the context of reduced Fur4 trafficking to the surface, as shown by localization defects of Fur4-mNG, and an unrelated transporter Mup1-GFP (**Figure 5F**). Collectively this example suggests that Uba4 and Met18 regulate expression of sufficient levels of Sec62, which are all required for proper trafficking of cargoes to the PM (**Figure 5G**).

**FIGURE 5:**
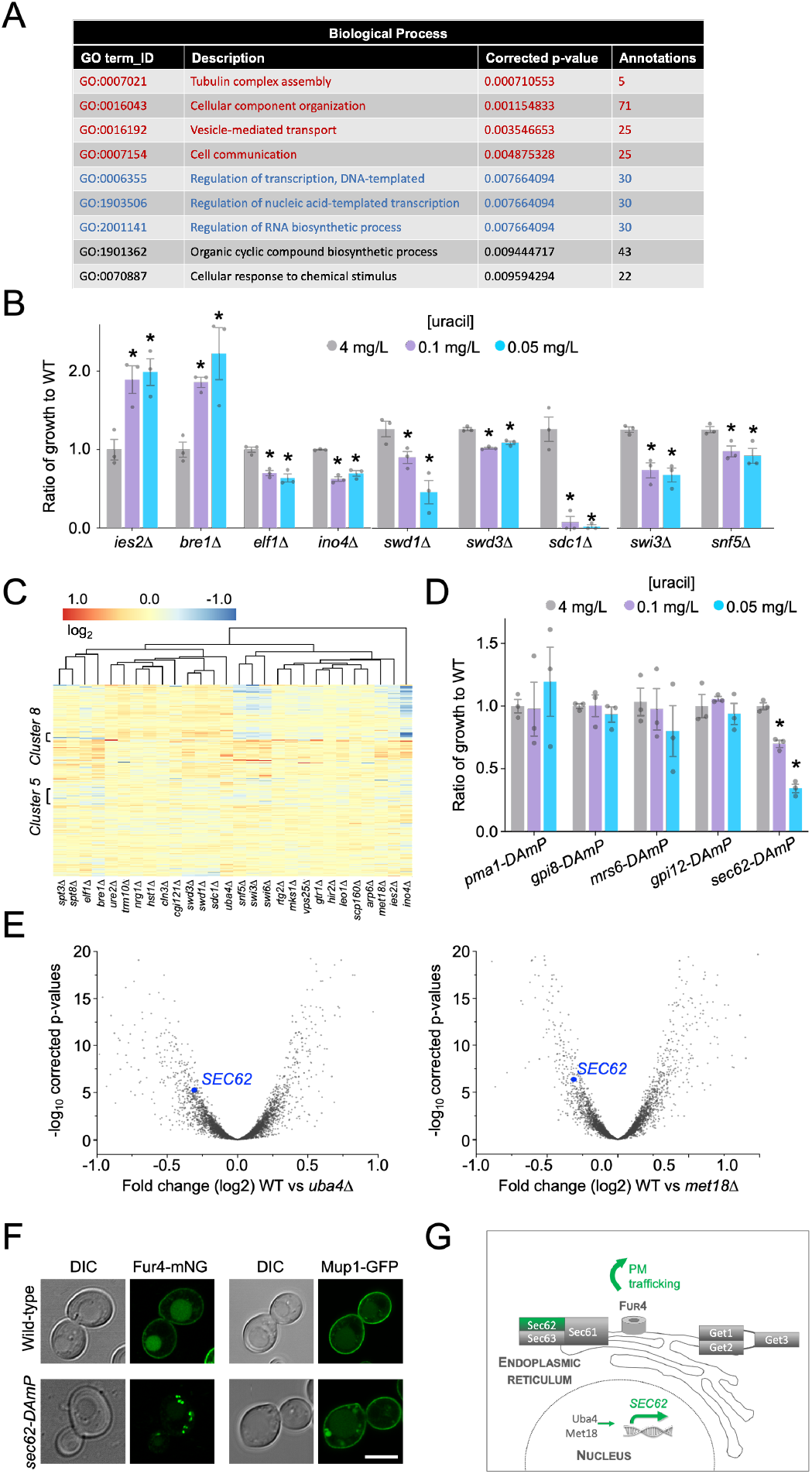
Trafficking screen enriched for transcriptional regulation. *A:* Details of Gene Ontology enrichments related to for biological processes for the 150 candidates identified in the screen. *B:* Ratio of growth for selected transcription factor mutants identified in the screen compared to wild-type cells at 4 mg/L, 0.1 mg/L and 0.05 mg/L uracil. Asterisks (*) indicate Student’s *t*-test comparisons p < 0.05. *C:* Hierarchical clustering and heat map of essential gene expression profiles in 28 different transcription factor null cells, coloured based on fold change. *D:* Ratio of growth for indicated mutant strains compared to wild-type (WT) cells grown on media containing 4 mg/L, 0.1 mg/L and 0.05 mg/ uracil. Asterisks (*) indicate Student’s *t*-test comparisons p < 0.0001. *E:* Volcano plots were constructed for log2 fold changes and their corresponding -log10 corrected p-values for genes in microarray analyses comparing wild-type cells to indicated mutants, with *SEC62* labelled (blue). *F:* Airyscan microscopy of Fur4-mNG or Mup1-GFP expressed from their respective endogenous promoters in wild-type and *sec62-DAmP* cells. Scale bar = 5 µm. *G:* Schematic diagram highlighting transcriptional and ER-associated factors from the screen that contribute to efficient trafficking of Fur4 to the surface.

### Uncharacterized factors are controlled at the transcriptional level

The uracil-scavenging screen identified ten uncharacterized candidates (**Figure 6A**). In an effort to understand whether these were also regulated by the TFs from the screen, we used a similar approach to cross-reference gene expression profiles of the TF-deletion strains against the 150 candidates identified in the screen. Again, hierarchical clustering revealed many gene profiles share signatures across different TF deletion experiments (**Figure 6B**). We were particularly intrigued by *ydr222w*Δ mutants defective in growth from the screen (**Figure 6A**) as *YDR222W* was greatly downregulated in a large number of the transcription factor null mutants, including the *swi3Δ, spt3*Δ and COMPASS complex mutants, but upregulated in other mutants not associated with low-uracil growth (**Figure 6C, Figure S4**). To test whether Ydr222w has a role in general surface protein trafficking, we expressed the distinct Mup1-GFP transporter in wild-type and *ydr222w*Δ mutants to reveal mislocalization phenotypes (**Figure 6D**) at both mid-log phase and late-log phase, the latter of which induces vacuolar sorting of Mup1-GFP^49,50^. Therefore, this bioinformatic approach helps prioritize factors, like the uncharacterized protein Ydr222w, for follow up testing and can be applied to other genetic screens, even retrospectively, that identify transcriptional regulators. To initiate functional categorisation of all the 150 factors, in particular where uncharacterised factors like Ydr222w might function, a secondary screen was performed based on the trafficking of α-factor through the secretory pathway to induce cell cycle arrest on a lawn of Mat A yeast, which are sensitised to arrest through a mutation in the gene that encodes the Bar1 mating factor protease (*bar1-1*). We hypothesised that mutants with defects in the secretory pathway leading to reduced alpha factor secretion would result in a reduction in the growth arrest response seen when Matα cells are spotted onto a lawn of MatA cells (**Figure 6E**).

**FIGURE 6:**
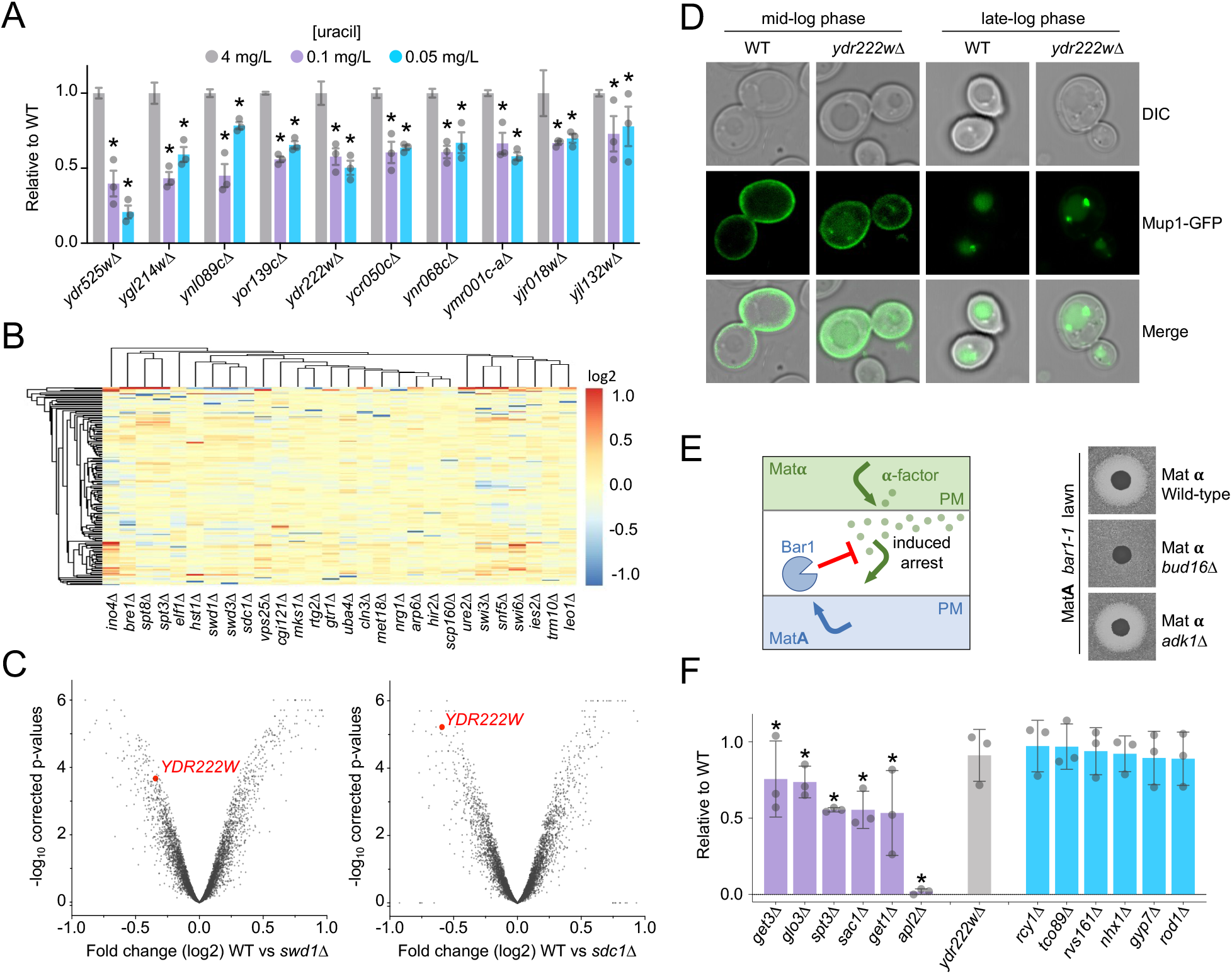
Previously uncharacterised proteins implicated in membrane trafficking. *A:* Ratio of growth compared to wild-type cells at 4 mg/L, 0.1 mg/L and 0.05 mg/L uracil for indicated uncharacterised mutants identified in the screen. Asterisks (*) indicate Student’s *t*-test comparisons p < 0.001. *B:* Hierarchical clustering of genes identified from the screen, plotting expression profiles as a heat map based on fold changes following deletion of 28 different transcriptional regulators. *C:* Volcano plots constructed for log2 fold changes and their corresponding p-values for genes in microarray analyses comparing wild-type cells to *swd1*Δ cells (left) and *sdc1*Δ cells (right). Value for *YDR222W* is shown in each plot (red). *D:* Wild-type and *ydr222w*Δ mutants were grown in serial dilution overnight and images at mid-log phase (OD600 = <1.0) or late log phase (OD600 = 2.0 + 2 hours) prior to confocal imaging. *E:* Schematic showing basis of α-factor induced secondary screen growing Mat**α** mutants from the primary screen on a lawn of *bar1-1* Mat *A* mutants (left), with representative examples shown (right). *F:* Quantification of growth inhibition surrounding spots of Mat**α** cells from screen described in *C*, shown relative to wild-type controls from same plate (n = 3). Asterisks (*) indicate Student’s *t*-test comparisons p < 0.01.

Conversely, mutants with no halo phenotype could act downstream of the PM, or indirectly, to explain uracil-scavenging phenotypes. We observed mutants to have a reduction in halo size (eg. *bud16*Δ) or a halo comparable to WT cells (eg. *adk1*Δ). We then performed growth arrest experiments in triplicate for all 150 screen candidates and found approximately half (76 / 150_ had statistically significant reduction in halo diameter (**Table S6**). Mutants of proteins well characterized to act in the early stages of the secretory pathway such as GET-complex nulls (discussed above) and *apl2*Δ^39,51^ were seen to display a significantly reduced halo phenotype. Whilst mutants of factors acting downstream of the secretory pathway in the endosomal system such as *rcy1*Δ and *nhx1*Δ^34,35^ were seen to display a halo phenotype not significantly different from WT (**Figure 6F**). Interestingly, uncharacterized mutant *ydr222w*Δ gave a halo type not significantly different to WT cells.

### Summary

In conclusion, we report a simple growth assay that indirectly reports on surface protein trafficking via nutrient transporter activity of uracil auxotroph yeast strains. The assay relies on comparison of growth efficiency of yeast cells on relatively high and low uracil media to infer the capacity of the Fur4 transporter to scavenge uracil required for growth. It is therefore cheap, simple and easy to perform at high throughput, as demonstrated by testing a haploid deletion library of over 5000 yeast strains. This genetic screen identified many novel candidates as potential Fur4 regulators and was particularly enriched for membrane trafficking and transcriptional machinery. By cross-referencing essential genes and factors identified from the screen, with genome-wide expression pattens in the majority of these transcriptional regulators, we were able to identify connections between TFs and the genes they regulate, both of which relate to uracil-scavenging. As an example, we document the essential gene *SEC62* and the uncharacterised gene *YDR222W*, as repressed in many of the TFs mutants identified from the screen. Although we cannot exclude the possibility that mutants have indirect effects on uracil scavenging, for example via the biosynthetic or metabolic processes. However, as membrane trafficking and transcription factors were significantly enriched, and many of the TFs can be functionally explained in the context of membrane trafficking, this suggests the bulk of mutants reported likely affect trafficking pathways used by Fur4. Furthermore, both factors prioritized by bioinformatics, Sec62 and Ydr222w, are shown to be required for proper sorting of fluorescently labelled cargoes. In addition to fluorescently labelled Fur4-mNG (**Figures 1, 2, 5**), and other surface cargoes affected by mutants from the screen, like Mup1-GFP and the G-protein coupled receptor Ste3 tagged with mCherry (**Figure S5**), we created mGFP tagged versions of both full length Fur4, or a mutant lacking its N-terminal 60 residues under copper inducible *CUP1* promoter, to allow temporal control of trafficking (**Figure S6**). We reasoned Fur4^ΔN^-mGFP would be a suitable reporter for secretory pathway mutants, as it cannot be endocytosed^17^, however localisation in *get1Δ, get2*Δ and *get3*Δ was indistinguishable from wild-type cells.

This might suggest the GET complex exhibits a distinct function in uracil-uptake. However, given the GET complex is known to be involved in secretory trafficking^39^ and not only were all 3 members identified from a blind screen of >5000 mutants, but *get1Δ get2*Δ and *get3*Δ cells were all defective in an assay reporting on secreted mating factor (0.53 ± 0.28, 0.80 ± 0.15, and 0.76 ± 0.24 respectively), we favour the explanation that the uracil-scavenging assay is sensitive enough to reveal a phenotype that is not apparent from steady state localization experiments. Therefore, we propose this uracil-scavenging assay, used in combination with fluorescently tagged cargoes, the mating factor secretion assay and bioinformatic approaches all documented herein serve as useful tools to study surface protein trafficking, in addition to the mutants identified and characterized, many of which are novel and evolutionarily conserved, that can inform future studies.

## METHODS

### Cell culture

Yeast cells were grown in rich yeast extract peptone dextrose (YPD) media (1% (w/v) yeast extract, 2% (w/v) peptone, 2% (w/v) D-glucose) or synthetic complete (SC) minimal media (0.675% (w/v) yeast nitrogen base without amino acids, 2% (w/v) D-glucose, plus the appropriate amino acid or base drop-outs for selection; (Formedium Norfolk, UK). Standard SC media contained 4 mg/L uracil and lower concentrations used, typically 0.1 mg/L and 0.05 mg/L for uracil stress conditions listed throughout. Cells were routinely cultured overnight to early/mid-log phase (OD600 < 1.0) prior to experimental procedures. Yeast strains used in this study are listed in **Table S7** and plasmids used are itemised in **Table S8**.

### Yeast growth assays

For the primary screen, 10μl culture of each mutant strain was grown in the well of a 96-well plate containing 150μl of YPD media with 250 µg/ml G418 overnight at 30°C in a humidified incubator. The bulk of YPD was then removed, followed by resuspension in water, and transfer of 10µl to a fresh 96-well plate containing 200µl sterile water. Dilutions were then mixed with a 96-pin replicator and pinned onto solid agar in 1,536 format using a ROTOR-HDA (Singer Instruments). Each mutant was pinned 16 times on solid media containing varying concentrations of uracil (4 mg/L or 0.1 mg/L or 0.05 mg/L) incubated at 30°C until sufficient growth was observed, followed by Phenobooth image capture (Singer Instruments) to record yeast growth. For follow up growth assays, the principle was the same but cells were cultured in 5ml serial dilutions to capture mid-log phase cells, which were then harvested with equivalent volumes to other strains, including a wild-type control, to be plated together. 6-step serial dilutions (10-fold) of each strain was generated in sterile water, followed by plating on SC plates containing of 4 mg/L, 0.1 mg/L and 0.05 mg/L uracil. Plates were then incubated at 30°C, growth recorded and intensity quantified after normalising for background (ImageJ; NIH).

### Confocal microscopy

Yeast cells expressing GFP and/or mCherry tagged proteins were grown to mid-log phase and then viewed in kill-buffer (50mM Tris pH 8.0, 10mM NaN3, 10mM NaF) or water (dH2O) at room temperature on Zeiss laser scanning confocal instruments (LSM710 or LSM880 equipped with Airyscan) with a Plan-Apochromat 63x/1.4 Differential Interference Contrast (DIC) objective lens. Images were captured using Zen Black Imaging software and modified using Image J (NIH). Yeast vacuoles were labelled with 1.6µM FM4-64 (ThermoFisher) in YPD media, followed by 3x washes with SC media and further growth in SC media for 1-hour prior to imaging. Where stated cells were fixed for imaging by spinning down mid-log cultures and washing with 100 mM potassium phosphate buffer (pH 8) at 7000rpm. Cells were resuspended and incubated for 10 minutes at room temperature in 4% paraformaldehyde (950µl K-Phos, 50 µl PFA) before spinning at 7000rpm and resuspending in 1 x PBS. Fixed cells were stained with Rhodamine-phalloidin (Phalloidin-594) and DAPI. When under the control of a copper inducible promoter

### α-factor induced arrest ‘halo’ assay

Wild-type and mutant Matα cells were grown to saturation overnight and then diluted back and grown for 4-6 hours in YPD media. Equivalent volumes of cells were harvested and spun down and brought up in 50 µl sterile water before spotting on low density lawns of MatA *bar1-1* cells created from mid-log phase cells on YPD plates. Plates were incubated at 30 degrees overnight. Regions of growth inhibition for mutants relative to 2x wild-type controls from the same plates were determined using ImageJ (NIH), with mean and standard error values calculated from three independent biological replicates.

### Gene Ontology analyses

All GO enrichments were acquired using the GO Term Finder v0.86^38^, using the default settings. The default background was used for all analyses except those for clusters derived from essential gene expression analysis, which instead used a background of essential genes listed in **Table S4**.

### Hierarchical clustering and gene expression analyses

Microarray data documenting changes in gene expression in mutant strains lacking transcriptional regulators compared to wild-type^46^ were assembled, representing 6123 genes. The data were read into R^52^ then processed using the dplyr v1.0.4^53^ and janitor v2.0.1^54^ packages to include only 28 strains lacking transcription factors identified in the uracil-scavenging genetic screen. For downstream bioinformatic analyses, two gene reference subsets were generated: the first included all 147 verified candidates from the screen, and the second included 1183 genes denoted as essential for viability. For all analyses, hierarchical clustering was performed using the pheatmap package v1.0.12^55^ using complete linkage. Elbow and silhouette analyses were performed using the factoextra package v1.0.7^56^ to determine the optimal number of clusters to guide further GO enrichment analyses. The Pearson correlation matrix of the whole genome expression data was produced with base R and visualised with the pheatmap package.

### String analysis

Interactome analysis of physical interactions was carried out using STRING software^57^, with a minimum required interaction score of 0.400 and FDR stringency of 5%. The full interaction network was clustered using k-means clustering (k=15).

### Statistical analysis

Statistical significance for experimental conditions were calculated using a student’s *t*-test/Bonferroni-Dunn method in GraphPad prism v8. Asterisks were used to denote significance on scatter plot histograms with p values documented in **Table S9**.

## Supporting information

Table S1_Uracil growth screen results and statistics

Table S2_Orthologues of screen candidates and associated human diseases

Table S3_GO enrichment annotations for biological processes

Table S4_Genes essential for viability used in bioinformatics

Table S5_List of genes from clusters of downregulation

Table S6_Mating factor induced growth arrest screen results and statistics

Table S7_Yeast Strains used in this study

Table S8_Plasmids used in this study

Table S9_Statistical significance tests

## ACKNOWLEDGEMENTS

We would like to thank Luke Mackinder, Jared Cartwright and the Protein Production Laboratory at the York Bioscience Technology Facility (BTF) for access and assistance with the Rotor HDA robotics, and the Imaging and Cytometry Core of the York BTF for technical support with microscopy. Thanks to Daphne Ezer for assistance with bioinformatics, and to Maya Schuldiner (Weizmann Institute, Israel) for providing DAmP-cassette integration yeast strains. This research was supported by a Sir Henry Dale Research Fellowship from the Wellcome Trust and the Royal Society 204636/Z/16/Z (CM) and a York Biology PhD studentship (KP).

## REFERENCES

1. Deshaies, R. J., Sanders, S. L., Feldheim, D. A. & Schekman, R. Assembly of yeast Sec proteins involved in translocation into the endoplasmic reticulum into a membrane-bound multisubunit complex. Nature 349, 806–808 (1991).

2. Stalder, D. & Gershlick, D. C. Direct trafficking pathways from the Golgi apparatus to the plasma membrane. Seminars in Cell and Developmental Biology 107, 112–125 (2020).

3. Laidlaw, K. M. E. & MacDonald, C. Endosomal trafficking of yeast membrane proteins. Biochemical Society Transactions 46, 1551–1558 (2018).

4. Lagunas, R. Sugar transport in Saccharomyces cerevisiae. FEMS Microbiol. Lett. 104, 229–242 (1993).

5. Nelson, N. Metal ion transporters and homeostasis. EMBO Journal 18, 4361–4371 (1999).

6. Perli, T., Wronska, A. K., Ortiz-Merino, R. A., Pronk, J. T. & Daran, J. M. Vitamin requirements and biosynthesis in Saccharomyces cerevisiae. Yeast 37, 283–304 (2020).

7. Grenson, M., Hou, C. & Crabeel, M. Multiplicity of the amino acid permeases in Saccharomyces cerevisiae. IV. Evidence for a general amino acid permease. J. Bacteriol. 103, 770–7 (1970).

8. Isnard, A. D., Thomas, D. & Surdin-Kerjan, Y. The study of methionine uptake in Saccharomyces cerevisiae reveals a new family of amino acid permeases. J. Mol. Biol. 262, 473–484 (1996).

9. Jund, R. & Lacroute, F. Genetic and Physiological Aspects of Resistance to 5-Fluoropyrimidines in Saccharomyces cerevisiae ’C-labeled pyrimidines and 32p were supplied by. (1970).

10. Babst, M. Eisosomes at the intersection of TORC1 and TORC2 regulation. Traffic 20, 543–551 (2019).

11. Bianchi, F., van’t Klooster, J. S., Ruiz, S. J. & Poolman, B. Regulation of Amino Acid Transport in Saccharomyces cerevisiae. Microbiol. Mol. Biol. Rev. 83, (2019).

12. Ljungdahl, P. O. & Daignan-Fornier, B. Regulation of amino acid, nucleotide, and phosphate metabolism in Saccharomyces cerevisiae. Genetics 190, 885–929 (2012).

13. Sardana, R. & Emr, S. D. Membrane Protein Quality Control Mechanisms in the Endo-Lysosome System. Trends in Cell Biology 31, 269–283 (2021).

14. Séron, K., Blondel, M. O., Haguenauer-Tsapis, R. & Volland, C. Uracil-induced down-regulation of the yeast uracil permease. Journal of Bacteriology 181, (1999).

15. Blondel, M.-O. et al. Direct Sorting of the Yeast Uracil Permease to the Endosomal System Is Controlled by Uracil Binding and Rsp5p-dependent Ubiquitylation. Mol. Biol. Cell 15, 883–895 (2004).

16. Hein, C., Springael J.-Y, Volland, C., Haguenauer-Tsapis, R. & André, B. NPI1, an essential yeast gene involved in induced degradation of Gap1 and Fur4 permeases, encodes the Rsp5 ubiquitin—protein ligase. Mol. Microbiol. 18, 77–87 (1995).

17. Keener, J. M. & Babst, M. Quality Control and Substrate-Dependent Downregulation of the Nutrient Transporter Fur4. Traffic 14, 412–427 (2013).

18. Moharir, A., Gay, L., Appadurai, D., Keener, J. & Babst, M. Eisosomes are metabolically regulated storage compartments for APC-type nutrient transporters. Mol. Biol. Cell 29, 2113–2127 (2018).

19. Appadurai, D. et al. Plasma membrane tension regulates eisosome structure and function. Mol. Biol. Cell 31, mbc.E19-04-0218 (2019).

20. Laidlaw, K. M. E. et al. A glucose-starvation response governs endocytic trafficking and eisosomal retention of surface cargoes in budding yeast. J. Cell Sci. 134, jcs.257733 (2021).

21. Boeke, J. D., La Croute, F. & Fink, G. R. A positive selection for mutants lacking orotidine-5₹-phosphate decarboxylase activity in yeast: 5-fluoro-orotic acid resistance. MGG Mol. Gen. Genet. 197, 345–346 (1984).

22. Winzeler, E. A. et al. Functional characterization of the S. cerevisiae genome by gene deletion and parallel analysis. Science (80-.). 285, 901–906 (1999).

23. Giaever, G. et al. Functional profiling of the Saccharomyces cerevisiae genome. Nature 418, 387–391 (2002).

24. Ghaemmaghami, S. et al. Global analysis of protein expression in yeast. Nature 425, 737–741 (2003).

25. Huh, W. K. et al. Global analysis of protein localization in budding yeast. Nature 425, 686–691 (2003).

26. Yofe, I. et al. One library to make them all: Streamlining the creation of yeast libraries via a SWAp-Tag strategy. Nat. Methods 13, 371–378 (2016).

27. Weill, U. et al. Genome-wide SWAp-Tag yeast libraries for proteome exploration. Nat. Methods 15, 617–622 (2018).

28. Arita, Y. et al. A genome-scale yeast library with inducible expression of individual genes. bioRxiv 2020.12.30.424776 (2021). doi:10.1101/2020.12.30.424776

29. Brachmann, C. B. et al. Designer deletion strains derived from Saccharomyces cerevisiae S288C: A useful set of strains and plasmids for PCR-mediated gene disruption and other applications. Yeast 14, (1998).

30. Novick, P., Field, C. & Schekman, R. Identification of 23 complementation groups required for post-translational events in the yeast secretory pathway. Cell 21, 205–215 (1980).

31. Franzusoff, A., Redding, K., Crosby, J., Fuller, R. S. & Schekman, R. Localization of components involved in protein transport and processing through the yeast Golgi apparatus. J. Cell Biol. 112, 27–37 (1991).

32. Babst, M., Katzmann, D. J., Estepa-Sabal, E. J., Meerloo, T. & Emr, S. D. ESCRT-III: An endosome-associated heterooligomeric protein complex required for MVB sorting. Dev. Cell 3, 271–282 (2002).

33. Teis, D., Saksena, S. & Emr, S. D. Ordered Assembly of the ESCRT-III Complex on Endosomes Is Required to Sequester Cargo during MVB Formation. Dev. Cell 15, 578–589 (2008).

34. Brett, C. L., Tukaye, D. N., Mukherjee, S. & Rao, R. The Yeast Endosomal Na (K)/H Exchanger Nhx1 Regulates Cellular pH to Control Vesicle Trafficking. Mol. Biol. Cell 16, 1396–1405 (2005).

35. Wiederkehr, A., Avaro, S., Prescianotto-Baschong, C., Haguenauer-Tsapis, R. & Riezman, H. The F-box protein Rcy1p is involved in endocytic membrane traffic and recycling out of an early endosome in Saccharomyces cerevisiae. J. Cell Biol. 149, 397–410 (2000).

36. MacDonald, C. & Piper, R. C. Genetic dissection of early endosomal recycling highlights a TORC1-independent role for Rag GTPases. J. Cell Biol. 216, 3275–3290 (2017).

37. Balakrishnan, R. et al. YeastMine-An integrated data warehouse for Saccharomyces cerevisiae data as a multipurpose tool-kit. Database 2012, (2012).

38. Cherry, J. M. et al. Saccharomyces Genome Database: The genomics resource of budding yeast. Nucleic Acids Res. 40, (2012).

39. Schuldiner, M. et al. The GET Complex Mediates Insertion of Tail-Anchored Proteins into the ER Membrane. Cell 134, 634–645 (2008).

40. Bulbarelli, A., Sprocati, T., Barberi, M., Pedrazzini, E. & Borgese, N. Trafficking of tail-anchored proteins: transport from the endoplasmic reticulum to the plasma membrane and sorting between surface domains in polarised epithelial cells. J. Cell Sci. 115, 1689–1702 (2002).

41. Geissler, S., Siegers, K. & Schiebel, E. A novel protein complex promoting formation of functional alpha-and gamma-tubulin. EMBO J. 17, 952–66 (1998).

42. Vainberg, I. E. et al. Prefoldin, a chaperone that delivers unfolded proteins to cytosolic chaperonin. Cell 93, 863– 73 (1998).

43. Millán-Zambrano, G. et al. The Prefoldin Complex Regulates Chromatin Dynamics during Transcription Elongation. PLoS Genet. 9, (2013).

44. Gerber, M. & Shilatifard, A. Transcriptional elongation by RNA polymerase II and histone methylation. Journal of Biological Chemistry 278, 26303–26306 (2003).

45. Zhou, H. et al. Snf5 and Swi3 subcomplex formation is required for SWI/SNF complex function in yeast. Biochem. Biophys. Res. Commun. 526, 934–940 (2020).

46. Kemmeren, P. et al. Large-scale genetic perturbations reveal regulatory networks and an abundance of gene-specific repressors. Cell 157, 740–752 (2014).

47. Breslow, D. K. et al. A comprehensive strategy enabling high-resolution functional analysis of the yeast genome. Nat. Methods 5, 711–718 (2008).

48. Schuldiner, M. et al. Exploration of the function and organization of the yeast early secretory pathway through an epistatic miniarray profile. Cell 123, 507–519 (2005).

49. MacDonald, C. et al. A Family of Tetraspans Organizes Cargo for Sorting into Multivesicular Bodies. Dev. Cell 33, 328–342 (2015).

50. Macdonald, C., Stringer, D. K. & Piper, R. C. Sna3 Is an Rsp5 Adaptor Protein that Relies on Ubiquitination for Its MVB Sorting. Traffic 13, 586–598 (2012).

51. Rad, M. R. et al. Saccharomyces cerevisiae Apl2p, a homologue of the mammalian clathrin AP β subunit, plays a role in clathrin-dependent Golgi functions. J. Cell Sci. 108, 1605–1615 (1995).

52. The R Foundation. R: The R Project for Statistical Computing, v4.0.4. (2021).

53. Wickham, H., François, R., Henry, L. & Müller, K. A Grammar of Data Manipulation [R package dplyr version 1.0.0]. (2020).

54. Firke, S. Simple Tools for Examining and Cleaning Dirty Data [R package janitor version 2.1.0]. (2021).

55. Kolde, R. CRAN - Package pheatmap. Cran.R-Project.Org (2019). Available at: https://cran.r-project.org/web/packages/pheatmap/index.html. (Accessed: 11th May 2021)

56. Alboukadel, Kassambara and Fabian, M. factoextra: Extract and Visualize the Results of Multivariate Data Analyses. R package. (2019).

57. Szklarczyk, D. et al. STRING v11: Protein-protein association networks with increased coverage, supporting functional discovery in genome-wide experimental datasets. Nucleic Acids Res. 47, D607–D613 (2019).

