## Supplemental Figures and Legends for "Fur4 mediated uracil-scavenging to screen for surface protein regulators"


- **Figure S1:** Methionine auxotroph trafficking mutants grow efficiently in low methionine media
- **Figure S2:** Interactome analysis of factors identified from screen
- **Figure S3:** Correlation matrix of responsive TF mutants of interest
- **Figure S4:** Expression of *YDR222W* in transcriptional mutants
- **Figure S5:** Fluorscently labelled surface cargoes are mislocalised in trafficking mutants
- **Figure S6:** Inducible constructs for mGFP tagged versions of Fur4


### SUPPLEMENTAL TABLES


- **Table S1:** Uracil growth screen results and statistics
- **Table S2:** Orthologues of screen candidates and associated human diseases
- **Table S3:** GO enrichment annotations for biological processes
- **Table S4:** Genes essential for viability used in bioinformatics
- **Table S5:** List of genes from clusters of downregulation
- **Table S6:** Mating factor induced growth arrest screen results and statistics
- **Table S7:** Yeast strains used in this study
- **Table S8:** Plasmids used in this study
- **Table S9:** Statistical significance tests


### SUPPLEMENTAL REFERENCES

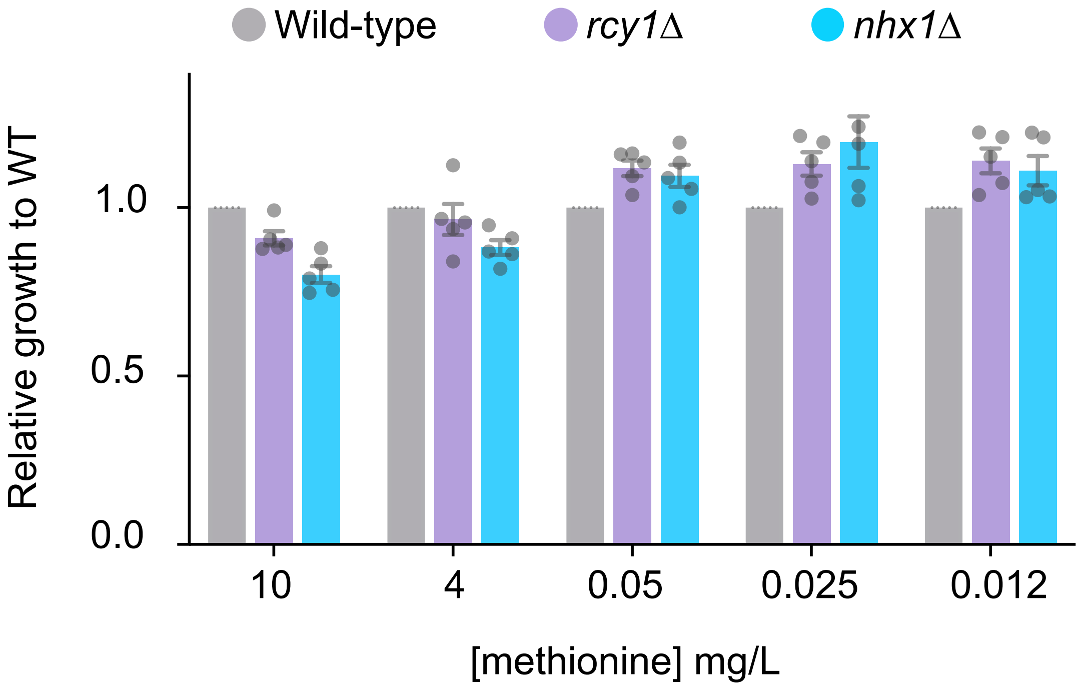


*Figure S1: Methionine auxotroph trafficking mutants grow efficiently in low methionine media*

Wild-type BY4741 Mat***A*** cells harbouring the *met15*Δ mutation that confers methionine auxotrophy, and two additional strains in this background additionally harbouring *rcy1*Δ (purple) or *nhx1*Δ (blue) mutations, were grown on SC media plates containing various indicated concentrations of methionine. There was no concentration of methionine that supported growth whilst also resulting in any significant difference in growth between strains.

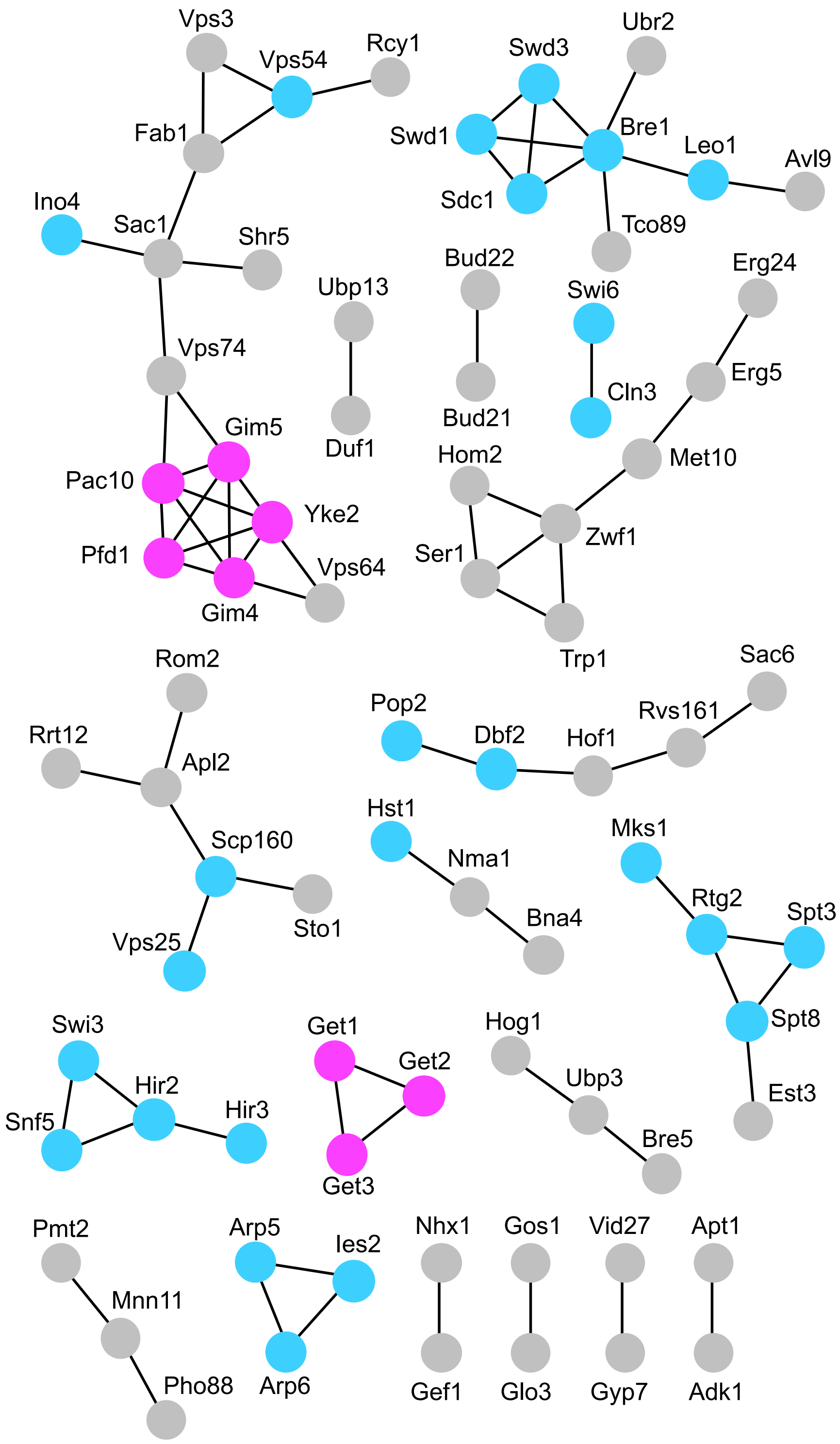


*Figure S2: Interactome analysis*

STRING pathway analysis for factors with known physical interactions identified in the low uracil-specific growth screen. Factors enriched for GO terms associated with cellular component (pink) and those that are included in bioinformatic assessment of gene expression upon their deletion (blue) are indicated. Orphan candidates with no interactions have been removed.

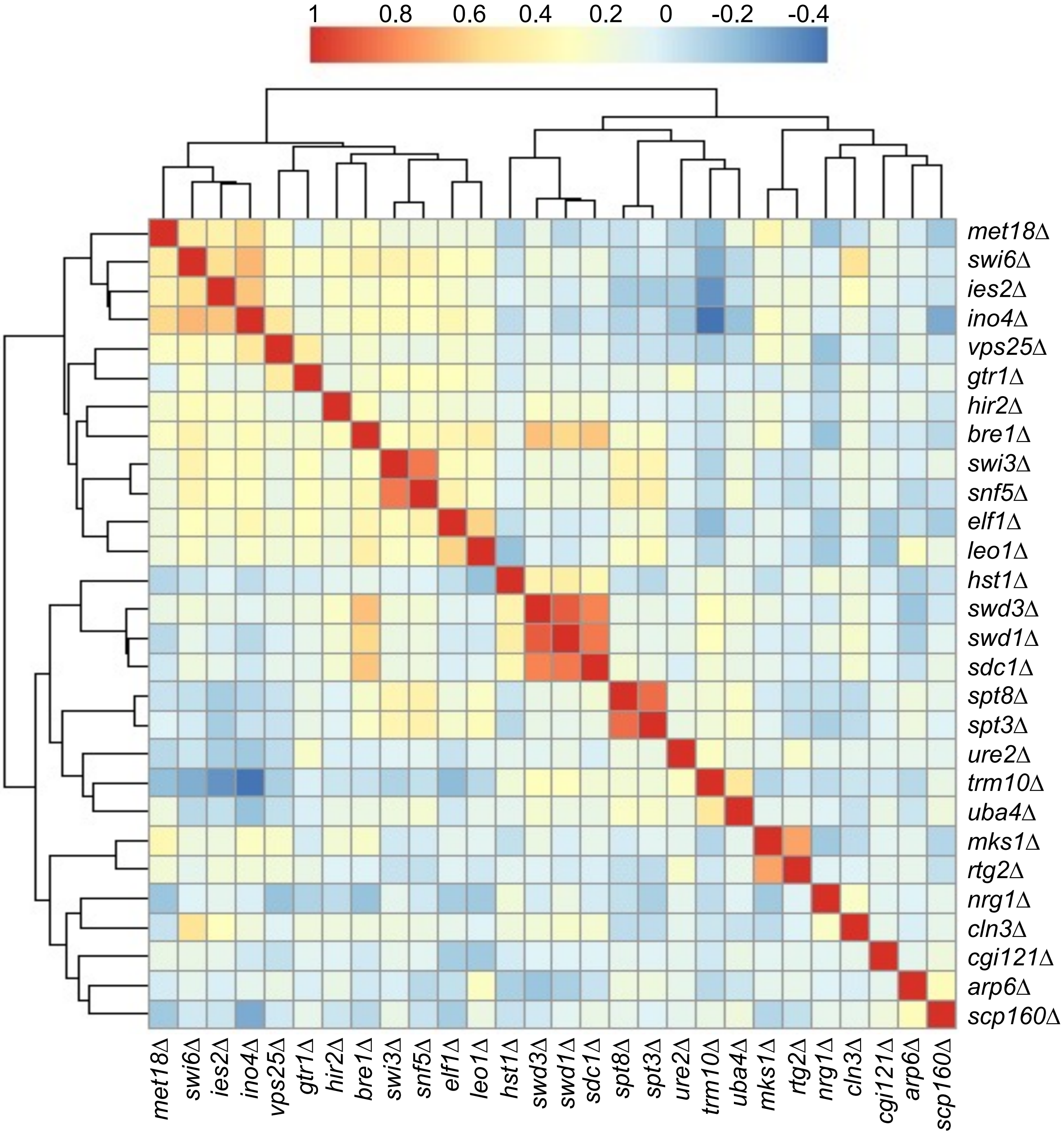


*Figure S3: Correlation matrix of responsive TF mutants of interest*

Correlation matrix of responsive mutants of interest, where 1 is perfect correlation as found from comparing identical mutants. Mutants of known interacting partners, for example Swd1, Swd3, Sdc1, exhibit high levels of correlation.

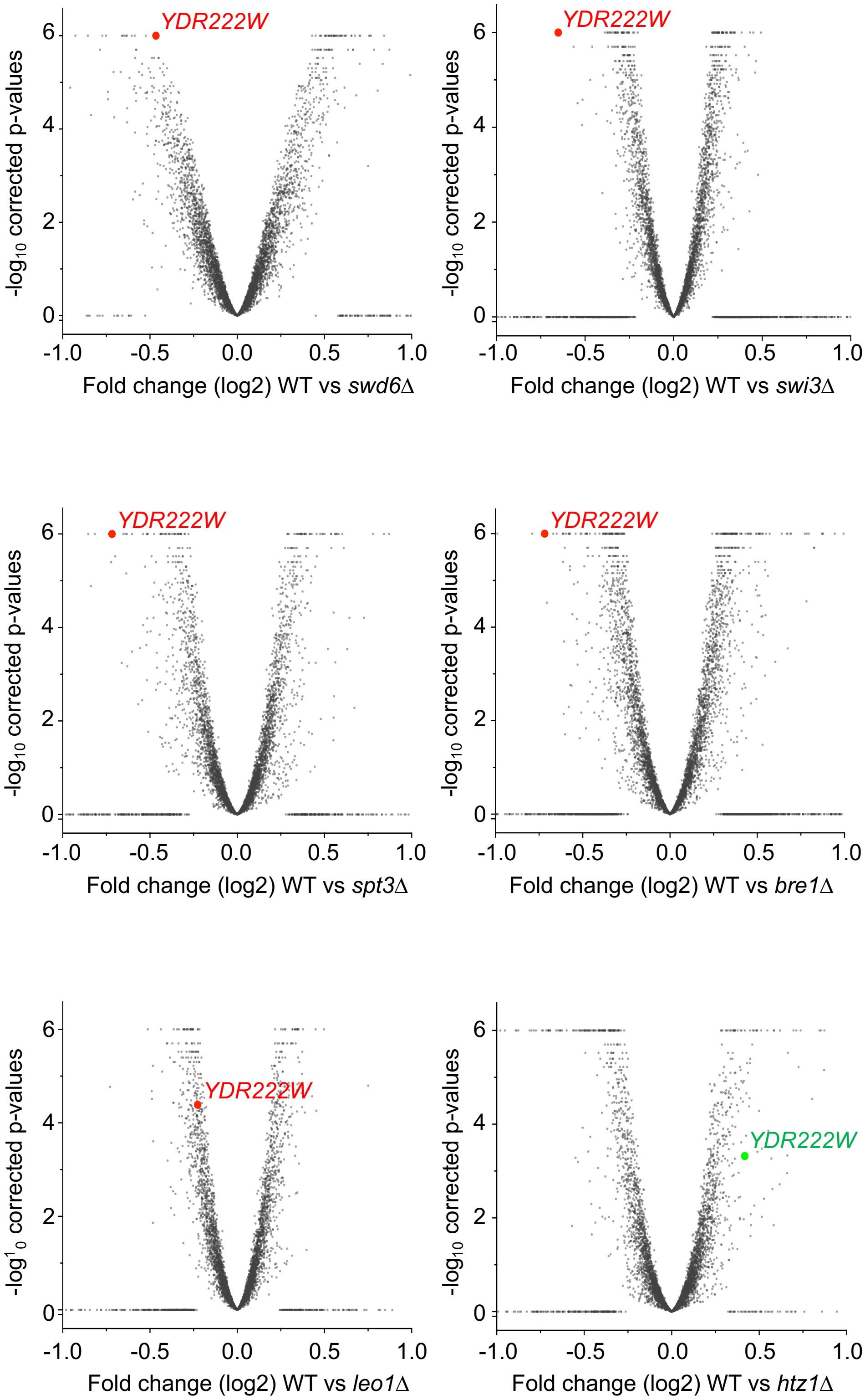


### *Figure S4: Expression of YDR222W in transcriptional mutants*

Volcano plots were constructed for $log_2$ fold changes and their corresponding p-values for genes in microarray analyses comparing wild-type cells to indicated mutants. Value for *YDR222W* expression is shown in each plot from mutants identified fro mthe screen (red) and an independent analysis of *htz1Δ* mutants that were not identified as Fur4-related from the screen (green).

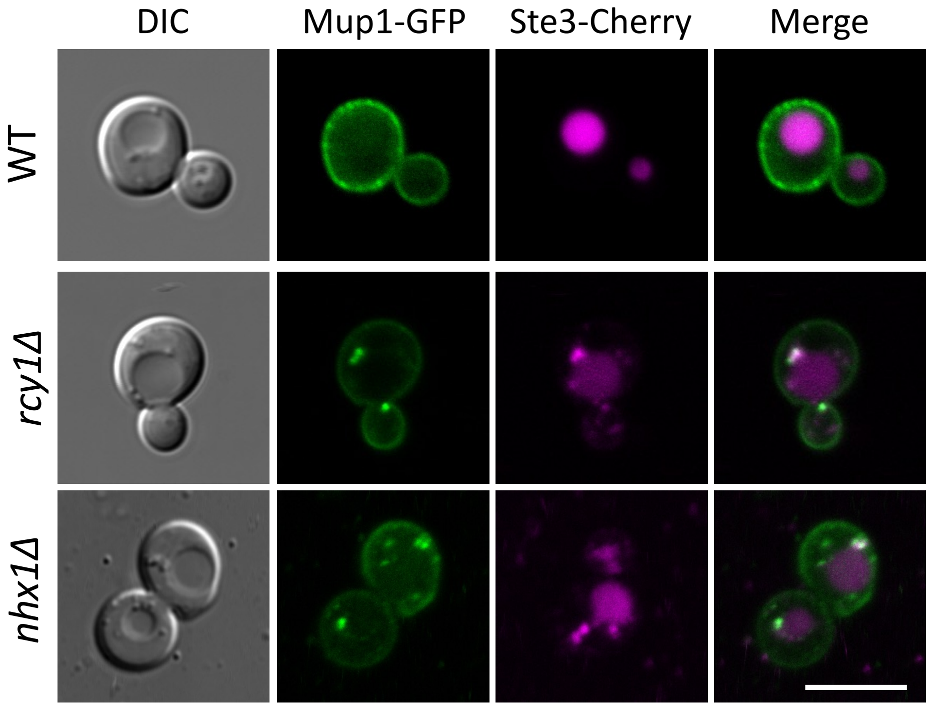


*Figure S5: Fluorscently labelled surface cargoes are mislocalised in trafficking mutants*

Indicated cells were co-transformed with plasmids expressing Mup1-GFP and Ste3-mCherry under control of endogenous promoters, grown to mid-log phase in selective media to retain plamids, prior to harvesting and imaging by confocal microscopy.

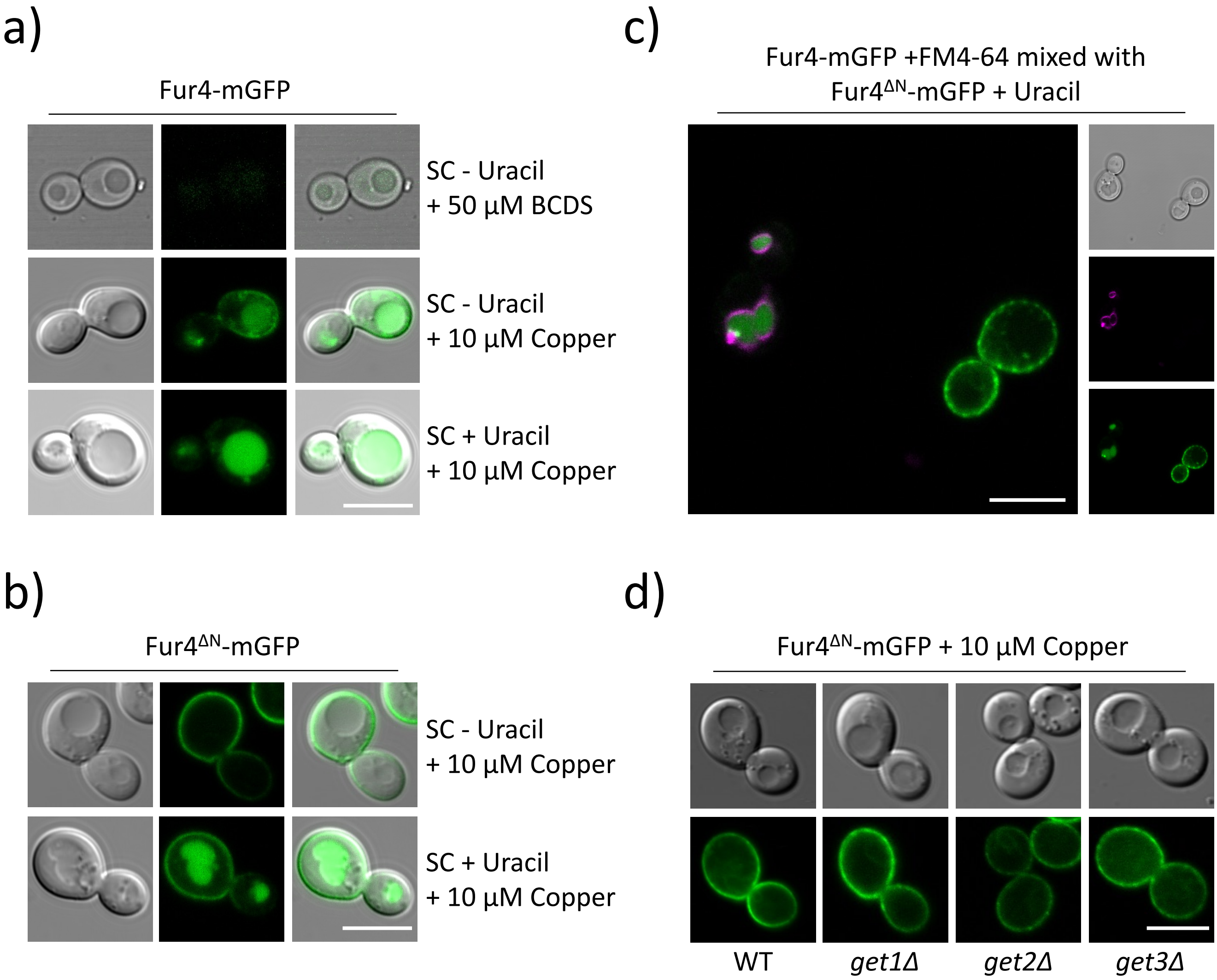


*Figure S6: Inducible constructs for mGFP tagged versions of Fur4*

***a)*** Wild-type cells transformed with a Fur4-mGFP expression plasmid under the control of the *CUP1* promoter were grown to mid-log phase in minimal media before addition of either 50 µM copper chelator BCDS or 10 µM copper chloride was added for 1 hour prior to imaging. ***b)*** Similarly, 10µM copper chloride was added to the media of log-phase grown cells transformed with a Fur4$^{\Delta N}$-mGFP expression plasmid, cells were imaged by confocal microscopy 1-hour after (lower). ***c)*** *Wild-type cells expressing Fur4-mGFP were labelled with FM4-64 for 1 hour, mixed with cells expressing* Fur4$^{\Delta N}$-mGFP *and co-cultured in the presence of 40 µg/ml uracil and* 10 µM copper chloride *prior to imaging.* ***d)*** Fur4$^{\Delta N}$-mGFP *was expressed in indicated strains followed by microscopy.* Scale bar, 5 µm

### SUPPLEMENTAL REFERENCES


1. Brachmann, C. B. *et al. Designer deletion strains derived from Saccharomyces cerevisiae S288C: A useful set of strains and plasmids for PCR-mediated gene disruption and other applications*. *Yeast* **14**, (1998).
2. Breslow, D. K. *et al.* A comprehensive strategy enabling high-resolution functional analysis of the yeast genome. *Nat. Methods* **5**, 711–718 (2008).
3. Ciejek, E. & Thorner, J. Recovery of S. cerevisiae a cells from G1 arrest by α factor pheromone requires endopeptidase action. *Cell* **18**, 623–635 (1979).
4. Laidlaw, K. M. E. *et al.* A glucose-starvation response governs endocytic trafficking and eisosomal retention of surface cargoes in budding yeast. *J. Cell Sci.* **134**, jcs.257733 (2021).
5. Winzeler, E. A. *et al.* Functional characterization of the S. cerevisiae genome by gene deletion and parallel analysis. *Science (80-. ).* **285**, 901–906 (1999).
6. MacDonald, C. *et al.* A Family of Tetraspans Organizes Cargo for Sorting into Multivesicular Bodies. *Dev. Cell* **33**, 328–342 (2015).
7. Giaever, G. *et al.* Functional profiling of the Saccharomyces cerevisiae genome. *Nature* **418**, 387–391 (2002).
8. Dowell, R. D. *et al.* Genotype to phenotype: A Complex problem. *Science* **328**, 469 (2010).
9. Rabitsch, K. P. *et al.* A screen for genes required for meiosis and spore formation based on whole-genome expression. *Curr. Biol.* **11**, 1001–1009 (2001).
10. Kastenmayer, J. P. *et al.* Functional genomics of genes with small open reading frames (sORFs) in S. cerevisiae. *Genome Res.* **16**, 365–373 (2006).
11. Bloom-Ackermann, Z. *et al.* A Comprehensive tRNA Deletion Library Unravels the Genetic Architecture of the tRNA Pool. *PLoS Genet.* **10**, (2014).
